# Descending and ascending signals that maintain rhythmic walking pattern in the cricket

**DOI:** 10.1101/2020.11.02.364422

**Authors:** Keisuke Naniwa, Hitoshi Aonuma

## Abstract

The cricket is one of the model animals used to investigate the neuronal mechanisms underlying adaptive locomotion. An intact cricket walks with a tripod gait, similar to other insects. The motor control center of the leg movements is located in the thoracic ganglia. In this study, we investigated the walking gait patterns of crickets whose ventral nerve cords were surgically cut to gain an understanding of how the descending signals from the head ganglia and ascending signals from the abdominal nervous system into the thoracic ganglia mediate the initiation and coordination of the walking gait pattern. Crickets whose paired connectives between the brain and subesophageal ganglion (SEG) were cut exhibited a tripod gait pattern. However, when one side of the connectives between the brain and SEG was cut, the crickets continued to turn in the opposite direction to the connective cut. Crickets whose paired connectives between the SEG and prothoracic ganglion were cut did not walk, whereas the crickets exhibited an ordinal tripod gait pattern when one side of the connectives was intact. Crickets whose paired connectives between the metathoracic ganglion and abdominal ganglia were cut initiated walking, although the gait was not a coordinated tripod pattern, whereas the crickets exhibited a tripod gait when one side of the connectives was intact. These results suggest that the brain plays an inhibitory role in initiating leg movements, and that both the descending signals from the head ganglia and the ascending signals from the abdominal nervous system are both important in initiating and coordinating insect walking gait patterns.

## 1 Introduction

One of the common issues between biologists and robotics scientists is revealing the mechanisms underlying adaptive locomotion in animals. It is generally believed that insects appeared on the earth roughly 400 million years ago and that there are approximately 1,000,000 insect species living on the earth. One of the reasons that insects have successfully evolved to spread across the earth may be that they developed adaptive locomotion. Locomotion is a crucial behavior for insects to obtain resources such as foods, territories, and mating partners. Revealing the neuronal mechanisms underlying locomotion in insects can aid in understanding the evolution of insect behaviors, as well as accelerate the development of novel design and control laws for legged robots.

Exploratory behavior to identify resources is initiated by the command signals generated in the brain. Thus, descending signals from the brain are necessary for the initiation of voluntary walking in both vertebrates and invertebrates (Kagaya and Takahata, 2011). External and internal signals are associated with the initiation of various behaviors. Chemical cues initiate exploratory behavior in insects because they are attracted by the chemical components of food and pheromones (Dethier, 1947). Auditory signals are another type of cue for attracting conspecific insects. For example, female crickets express phonotaxis to the calling song stridulated by males (Alexander, 1961). Internal signals also function to initiate behaviors. Starvation and thirst can increase the motivation to initiate exploratory behavior for food and water, indicating that food digestion and the excretion system are associated with initiating behaviors in insects.

Insects are hexapod animals and most of them exhibit a tripod gait pattern, whereby the foreleg and hindleg on one side move in synchrony with the midleg on the other side (Wilson, 1966). Descending signals are important for initiating walking in insects (Strausfeld, 1999) (Emanuel et al., 2020). The local centers of the leg movements lie within the thoracic ganglia, where oscillatory neuronal activities, which are known as central pattern generators (CPGs), contribute to rhythmic leg movements (Borgmann et al., 2009). Descending information from the brain into the thoracic ganglia is necessary to coordinate the movement of the legs (Heinrich, 2002). The subesophageal ganglion (SEG) plays a crucial role in walking (Knebel et al., 2018a). However, our previous study demonstrated that headless crickets do not exhibit voluntary walking, except following defecation (Naniwa et al., 2019). After-defecation walking is initiated by ascending signals from the terminal abdominal ganglion. The gait of after-defecation walking in the headless cricket is not a coordinated tripod pattern.

In this study, we aimed to determine how the ascending signals from the abdominal nervous system influence the coordinated walking gait pattern. To investigate this issue, we surgically cut the connectives of the ventral nerve cord at different positions and analyzed the walking gait pattern of the field cricket. To determine the roles of the brain in initiating and regulating the walking gait, either the paired connectives or one side of the connectives between the brain and SEG were cut. To investigate the roles of the SEG, either the paired connectives or one side of the connectives between the SEG and prothoracic ganglion were cut. Furthermore, to investigate the roles of the ascending signals from the abdominal nervous system, either the paired connectives or one side of the connectives between the metathoracic ganglion and first free abdominal ganglion were cut. Based on these results, we demonstrated that both the descending signals and the ascending signals into the thoracic ganglia play an important role in maintaining a coordinated walking pattern.

## 2 Materials and methods

### 2.1 Animals

The cricket *Gryllus bimaculatus* (De Geer) used in this study were raised in a laboratory colony. They were reared on a 14 h:10 h light and dark cycle (lights on at 6:00 h) at 28 ± 2 °C. They were fed a diet of insect food (Sankyo Lab, Tokyo, Japan) and water ad libitum. Adult male crickets that had molted within two weeks before the experiments were randomly selected for use in this study.

### 2.2 Behavior experiments

The crickets used were randomly selected from the colony. A cricket was placed on a handmade treadmill to observe its walking pattern. The treadmill was composed of a Styrofoam sphere (*ϕ* 150 mm) that hovered over a stream of air flowing beneath it. Each cricket was anesthetized with CO_2_ gas before it was placed on the treadmill. A steel rod (*ϕ* 100 μm) was attached to the thorax of the cricket using dental wax (Shofu, Kyoto, Japan). The rod was inserted into a plastic tube (*ϕ* 500 μm) that was fixed to a manipulator, by means of which the cricket was placed in the exact desired position on the Styrofoam sphere. A cricket on the treadmill could walk as well as change its orientation and ground clearance freely.

To investigate the roles of either the ascending or descending signals into the thoracic ganglia, where the premotor signals for locomotion are generated, the intersegmental connectives between the brain and SEG, between the SEG and prothoracic ganglion, and between the metathoracic ganglion and abdominal ganglia were cut using a razor blade. The locomotion patterns of the crickets were observed and recorded using a high-speed camera (HAS-L1, DITECT, Japan, 800 × 640 pixels, 300 fps). The images were saved as sequential JPEG files on a Windows PC for subsequent analysis.

### 2.3 Data analysis

To analyze and evaluate the leg movement patterns, we drew polar histograms (Naniwa et al., 2020), in which we focused on the leg movement direction. In brief, we defined the power stroke as the thrust produced when the angle between the femur and tibia increased in the case of the hindleg, or when the angle between the femur and trunk increased in the case of the foreleg and midleg. During the recovery stroke, the angle between the femur and tibia decreased for the hindleg or the angle between the femur and trunk decreased for the foreleg and midleg. The stroke mode was obtained manually from the video data. The condition of each leg in a frame was compared to those of the adjacent frames to determine whether it was a power or recovery stroke.

In the definition of the phase for each leg, *t* is a certain time and *t_n_* is the start time of the power stroke directly before the *n*th step of the leg of interest.

The phase *ϕ* at time *t* is defined as

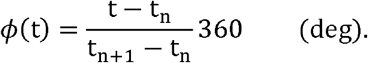

Therefore, the leg phase is defined as the period between the beginning of two consecutive power strokes. In this case, *ϕ*_object_, *ϕ*_subject_ are The phases of an arbitrary leg, where the subscripts object and subject indicate the leg positions (e.g., LF, RM).

The leg phase difference of the subject leg relative to the object leg at time *t* is expressed as

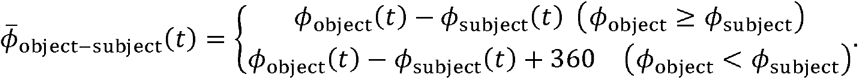

This method aims to provide an intuitive and a precise representation of the rhythmic pattern corresponding to the variations in the cricket legs owing to movement. Therefore, even in a polar representation, in which the area represents the ratio of frequencies, the height of a bar is the value of the square root of the frequency that it represents. As a result, the total area of the bar is 1 in a polar histogram (Nemec, 1988). The phase difference between the legs can be calculated for each frame. In an ideal tripod, the leg phase difference between adjacent legs (e. g., LF and RF or LF and LM) is always 180°.

In the polar histogram, the phase mean *Φ* is calculated as:

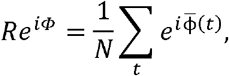

where *R* is the mean resultant length of each histogram, *N* is the total amount of sample data, and *i* is an imaginary number.

The circumferential dispersion *s* and circumferential standard deviation *ν* of the circumferential data are defined as follows:

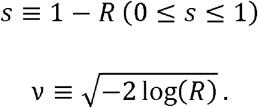

The rank statistics of the measured circumference data 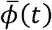, sorted in ascending order in the range of 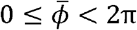, are represented by 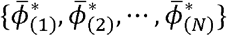.

In this case, the empirical distribution function *S*(*ϕ*) can be expressed as

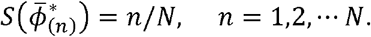

The variations in the phase differences between legs could be intuitively understood by comparing the shapes of the empirical distribution functions. In this paper, the empirical distribution function of the leg phase difference between the midlegs in each experiment is illustrated as a representative example.

## 3 Results

The intact crickets exhibited a tripod gait pattern during walking on the treadmill (Fig. 1, Sup1 movie). The polar histogram indicates the phase difference between two of the six legs. The phase difference between the left and right forelegs occurred in an almost anti-phase manner. The mean of the foreleg phase difference *Φ_LF–RF_* was 185°, with a standard deviation *ν_LF–RF_* of 39.6°. The mean vector length *R_LF–RF_* was 0.79. Similarly, the left and right midlegs moved in an anti-phase manner. The mean of the midleg phase difference *Φ_LM-RM_* was 164°, with a standard deviation *ν_LM–RM_* of 47.6°. The mean vector length *R_LM–RM_* was 0.71. The left and right hindlegs also moved in an anti-phase manner. The mean of the midleg phase difference *Φ_LH–RH_* was 180°, with a standard deviation *ν_LH–RH_* of 28.9°. The mean vector length *R_LH–RH_* was 0. 88. However, the foreleg and midleg on the same side moved in an almost anti-phase manner (*Φ_LF–LM_*: 212°, *ν_LF–LM_*: 42.2°, *R_LF–LM_*: 0.76, *Φ_RF–RM_*: 194°, *ν_RF–RM_*: 35.1°, *R_RF–RM_*: 0.83), and the foreleg and hindleg on the same side moved slightly later than in-phase 62.1°, *R_LF–LM_*: 0.56, *Φ_RF–RH_*: 71.7°, *v_RF–RH_*: 57.9°, *R_RF–RH_*: 0.60). These results suggest that the legs did not maintain a perfectly coordinated relationship with one another during the tripod gait in the intact crickets.

**Figure 1.**
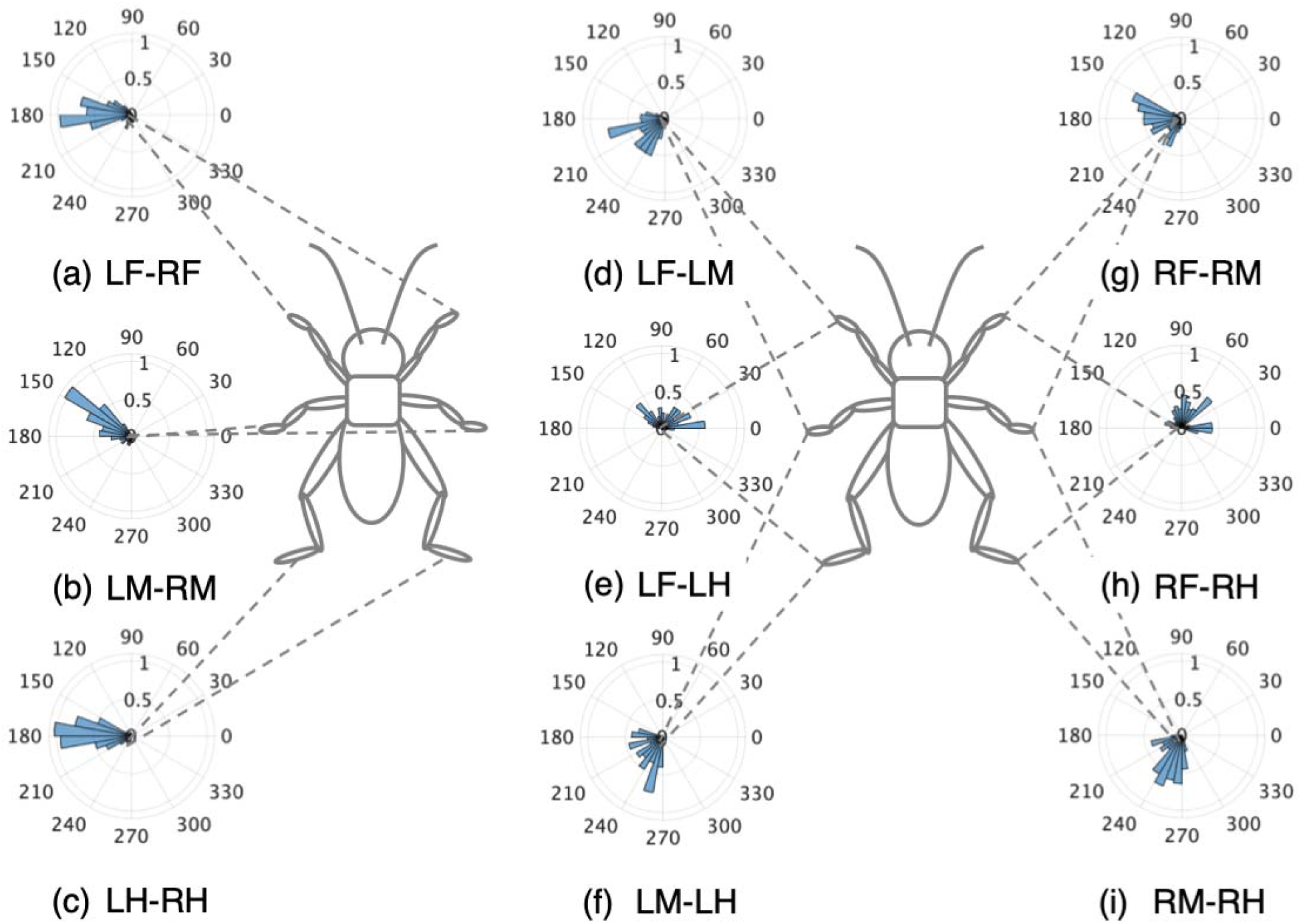
Walking gait patterns of intact crickets. Polar histograms indicating the phase differences between two legs in the intact crickets (N = 5), where the radial axis is the probability that the intact crickets exhibited a tripod gait pattern on the treadmill, (a) Phase difference between left and right forelegs, (b) Phase difference between left and right middle legs, (c) Phase difference between left and right hindlegs, (d) Phase difference between left foreleg and midleg. (e) Phase difference between left foreleg and hindleg, (f) Phase difference between left midleg and hindleg, (g) Phase difference between right foreleg and midleg. (h) Phase difference between right foreleg and hindleg, (i) Phase difference between left midleg and hindleg. LF: left foreleg, RF: right foreleg, LM: left midleg, RM: right midleg, LH: left hindleg, and RH: right hindleg.

To investigate the manner in which the ordinary tripod gait pattern is regulated by descending signals from the brain or ascending signals from the abdominal nervous system, the connectives of the ventral nerve cord were surgically disconnected. The central nervous system of insects has a symmetric structure. The brain (protocerebrum, deutocerebrum, and tritocerebrum) is joined by paired nerve connectives to the SEG, which is, in turn, linked to the thoracic and abdominal ganglia by paired connectives.

### 3.1 Disconnection of connectives between brain and SEG

The disconnection of the paired connectives between the brain and SEG did not change the walking gait pattern of the test crickets, which walked on the treadmill with a tripod gait (Fig. 2(A), Sup2 movie). The test crickets did not respond to tactile stimuli on the antennae, although they responded to tactile stimuli on the cercus while walking. This indicates that the descending signals from the brain into the SEG were shut down. The crickets mainly walked straight forward and did not turn voluntarily. The phase difference between the left and right midlegs occurred in an anti-phase manner (Fig. 2(A)(b)). In the intact crickets, the mean midleg phase difference *Φ_LM–RM_* was 164°, with a standard deviation *v_LM–RM_* of 47.6°. The mean vector length *R_RM–RM_* was 0.71. In contrast, in the test crickets, the mean of the midleg phase difference *Φ_LM–RM_* was 193°, with a standard deviation *ν_LM–RM_* of 55.5°. The mean vector length *R_LM–RM_* was 0.63. The shape of the empirical distribution function of the midlegs of the test crickets was similar to that of the intact crickets (Fig. 2(B)). This indicates that the test crickets walked with a gait similar to that of the intact crickets; that is, a tripod gait.

**Figure 2.**
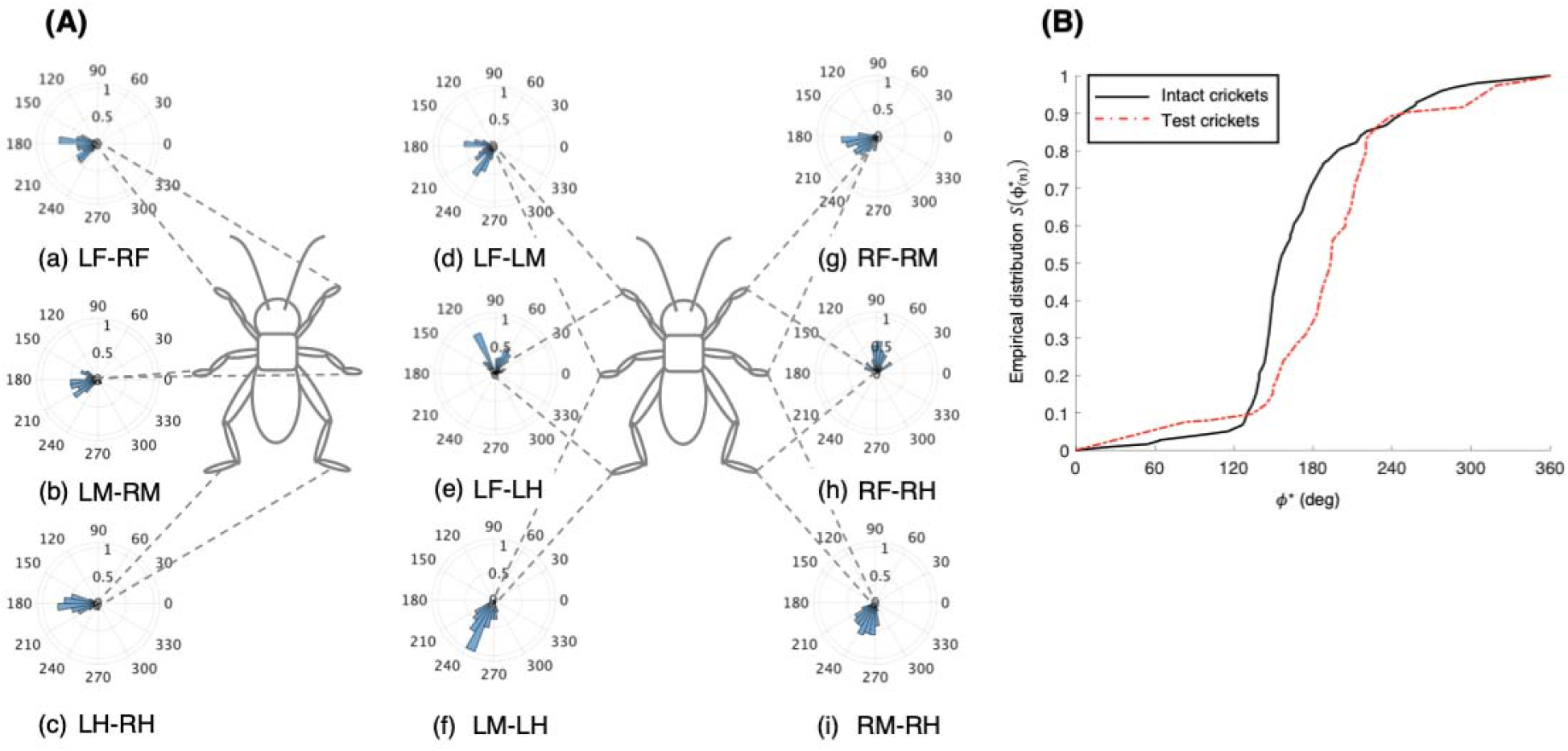
Walking gait pattern of crickets in which paired nerve connectives between brain and SEG were cut. **(A)** Polar histograms showing phase differences between two legs in test crickets (N = 5). The polar histograms demonstrate that the test crickets exhibited a tripod gait pattern on the treadmill, (a) to (i) Phase differences between pairs of legs. **(B)** Comparison of empirical distribution function of leg phase differences between left and right legs *Φ_LM–RM_* The black line indicates the empirical distribution function of the intact crickets and the red line indicates that of the crickets in which the pair of connectives between the brain and SEG was cut. LF: left foreleg, RF: right foreleg, LM: left midleg, RM: right midleg, LH: left hindleg, and RH: right hindleg.

However, the crickets in which only the left side of the connectives between the brain and SEG was cut did not walk straight forward, but continued to turn in the right direction (Sup3 movie). Their gaits did not exhibit not an ordinary tripod pattern (Fig. 3). The polar histogram of these test crickets indicates that the phase differences between the left and right legs were not consistent (Fig. 3(A)). In the test crickets, the mean of the midleg phase difference *Φ_LM–RM_* was 187°, with a standard deviation *ν_LM–RM_* of 131°. The mean vector length *R_RM–RM_* was 0.07. The shape of the empirical distribution function of the midlegs in the test crickets was different from that of the intact crickets (Fig. 3(B)). This analysis also demonstrates that the walking pattern was far from the ordinary tripod gait (Fig. 3(C)). The gait chart diagram of the test crickets reveals that the duration of the left leg movements appeared to be rhythmic, similar to that of the intact crickets. The duration of the right legs touching the floor was much longer than that of the left legs. This indicates that the left legs moved more than the right legs, making the cricket continue to turn right. Similarly, when the right side of the connectives between the brain and SEG was cut, the test crickets continued to turn left and did not exhibit a tripod walking gait pattern.

**Figure 3.**
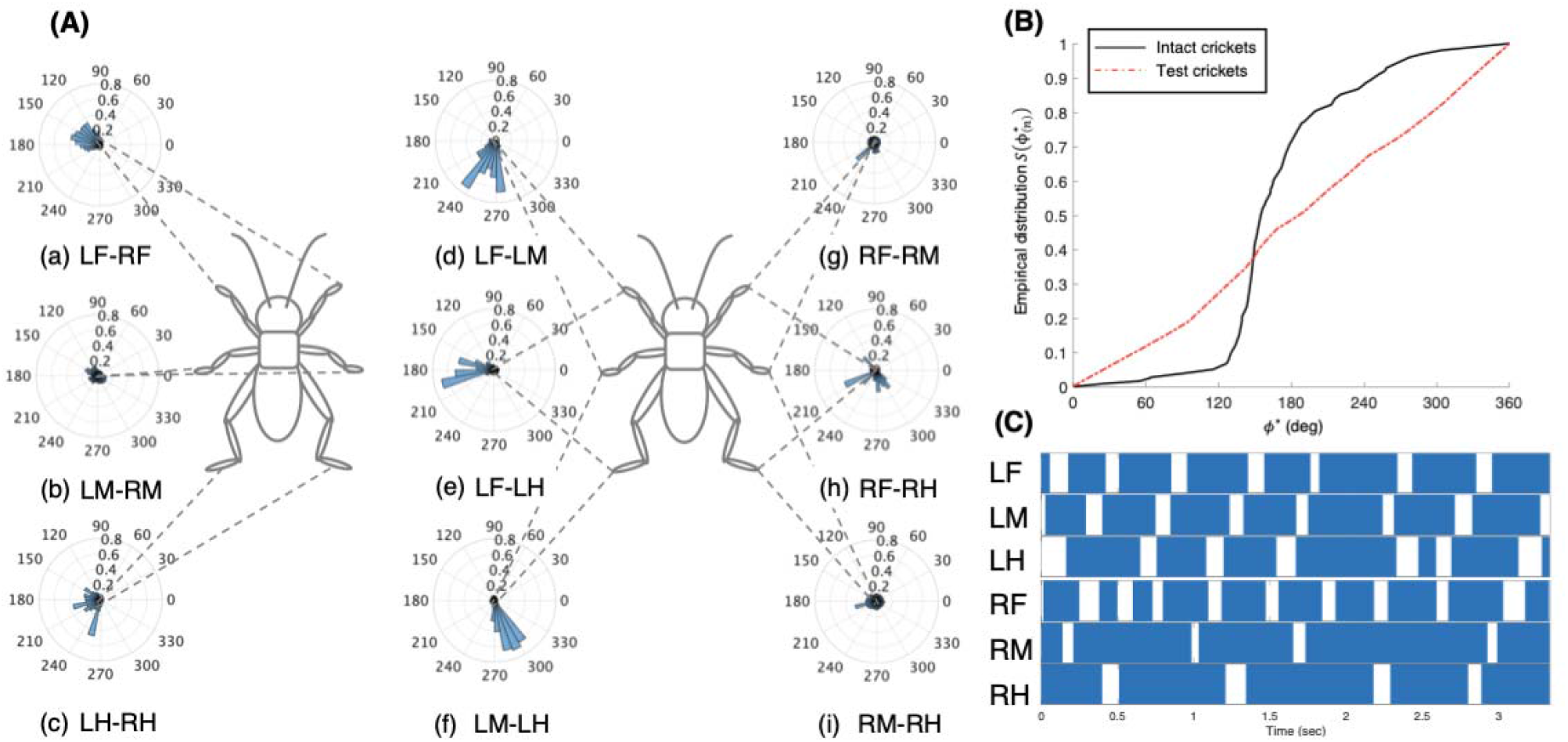
Walking gait patterns of crickets in which left side of nerve connectives between brain and SEG was cut. **(A)** Polar histograms showing phase differences between two legs in test crickets (N = 5). The test crickets continued to turn in the right direction. The polar histograms demonstrate that the walking pattern was not a tripod gait, (a) to (i) Phase differences between pairs of legs. **(B)** Comparison of empirical distribution function of leg phase differences between left and right legs *Φ_LM–RM_*. The black line indicates the empirical distribution function of the intact crickets and the red line indicates that of the test crickets. **(C)** Gait chart diagram of test cricket. The filled part indicates the duration of the power stroke period and the blank part indicates the duration of the recovery stroke. This also demonstrates that the walking pattern was not a tripod gait in the test cricket. LF: left foreleg, RF: right foreleg, LM: left midleg, RM: right midleg, LH: left hindleg, and RH: right hindleg.

### 3.2 Disconnection of connectives between SEG and prothoracic ganglion

To investigate the role of the SEG, the paired connectives in the crickets were surgically cut. The behavior of these test crickets was the same as those of the headless crickets previously reported (Naniwa et al., 2019). The test crickets did not walk, except during defecation. Therefore, the gait chart diagrams indicate that all legs of the crickets were always on the ground (Fig. 4). However, if only one of the connectives between the SEG and prothoracic ganglion was cut, the crickets were able to behave like intact crickets. The test crickets in which the left–side connective between the SEG and prothoracic ganglion was cut could walk with a tripod gait (Fig. 5). The phase differences between the left and right legs occurred in an anti–phase manner (Figs. 5(A) (a) to (c)). The foreleg and midleg of the same side moved in an anti–phase manner, whereas the foreleg and hindleg of the same side moved in an in-phase manner (Figs. 5(A)(d) to (i)). In the test crickets, the mean of the midleg phase difference *Φ_LM–RM_* was 165°, with a standard deviation *ν_LM–RM_* of 37.3°. The mean vector length *R_LM–RM_* was 0.81. The shape of the empirical distribution function of the pair of midlegs of the test crickets was similar to that of the intact crickets (Fig. 5(B)). We also examined the behavior when only the right–side connective between the SEG and prothoracic ganglion was cut. The results were quite similar to those of the crickets with the left–side connective cut (N = 5).

**Figure 4.**
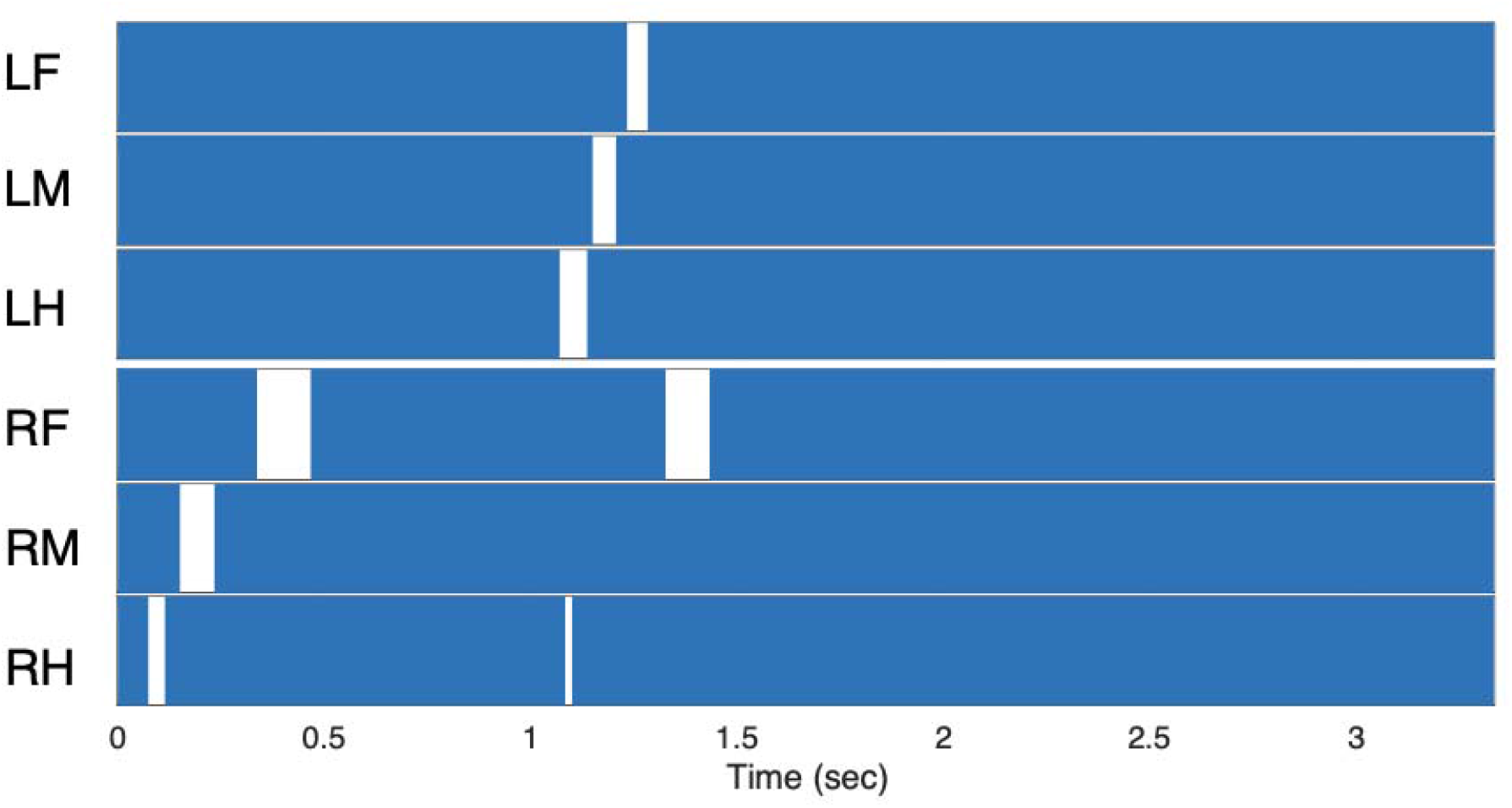
Gait chart diagram of cricket in which paired nerve connectives between SEG and prothoracic ganglion were cut. The filled part indicates that the tip of the legs touched the floor, demonstrating that the cricket did not walk on the treadmill. LF: left foreleg, RF: right foreleg, LM: left midleg, RM: right midleg, LH: left hindleg, and RH: right hindleg.

**Figure 5.**
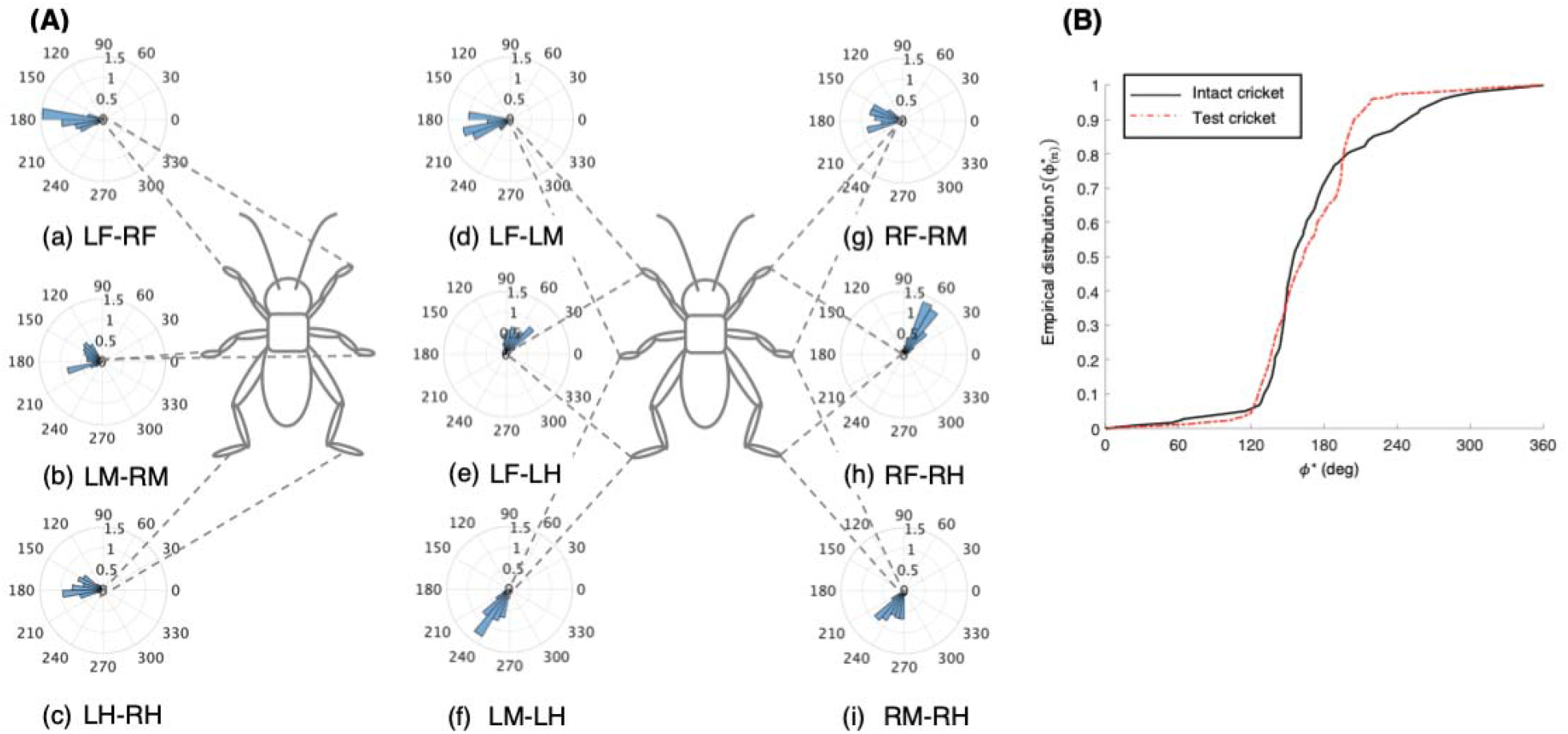
Walking gait patterns of crickets in which left side of nerve connectives between SEG and prothoracic ganglion was cut. **(A)** Polar histograms showing phase differences between two legs in test crickets (N = 5). The polar histograms demonstrate that the test crickets exhibited a tripod gait pattern on the treadmill, (a) to (i) Phase differences between pairs of legs. **(B)** Comparison of empirical distribution function of leg phase differences between left and right legs *Φ_LM–RM_*. The black line indicates the empirical distribution function of the intact crickets and the red line indicates that of the crickets in which only the left side of the nerve connectives between the metathoracic ganglion and abdominal ganglia was cut. LF: left foreleg, RF: right foreleg, LM: left midleg, RM: right midleg, LH: left hindleg, and RH: right hindleg.

The crickets in which the left–side connectives between both the brain and SEG, and the SEG and prothoracic ganglion were cut did not walk straight forward, but continued to turn in the right direction (N = 3, Sup4 movie). The gaits in these test crickets did not exhibit an ordinary tripod pattern (Fig. 6). The polar histogram of the test crickets indicates that the phase differences between the left and right legs were not consistent (Fig. 6(A)). In the test crickets, the mean of the midleg phase difference *Φ_LM–RM_* was 109°, with a standard deviation *ν_LM–RM_* of 108°. The mean vector length *R_LM–RM_* was 0.17. The shape of the empirical distribution function of the midlegs of the test crickets was different from that of the intact crickets (Fig. 6(B)). The gait chart diagram of the test crickets demonstrates that the duration of the left leg movements appeared to be rhythmic, as in the intact crickets (Fig. 6(C)). The duration of the right legs touching the floor was much longer than that of the left legs. We also investigated the behavior of the crickets in which the right–side connectives between both the brain and SEG, and between the SEG and prothoracic ganglion were cut (N = 3). These crickets continued to turn in the left direction and did not exhibit a tripod gait.

**Figure 6.**
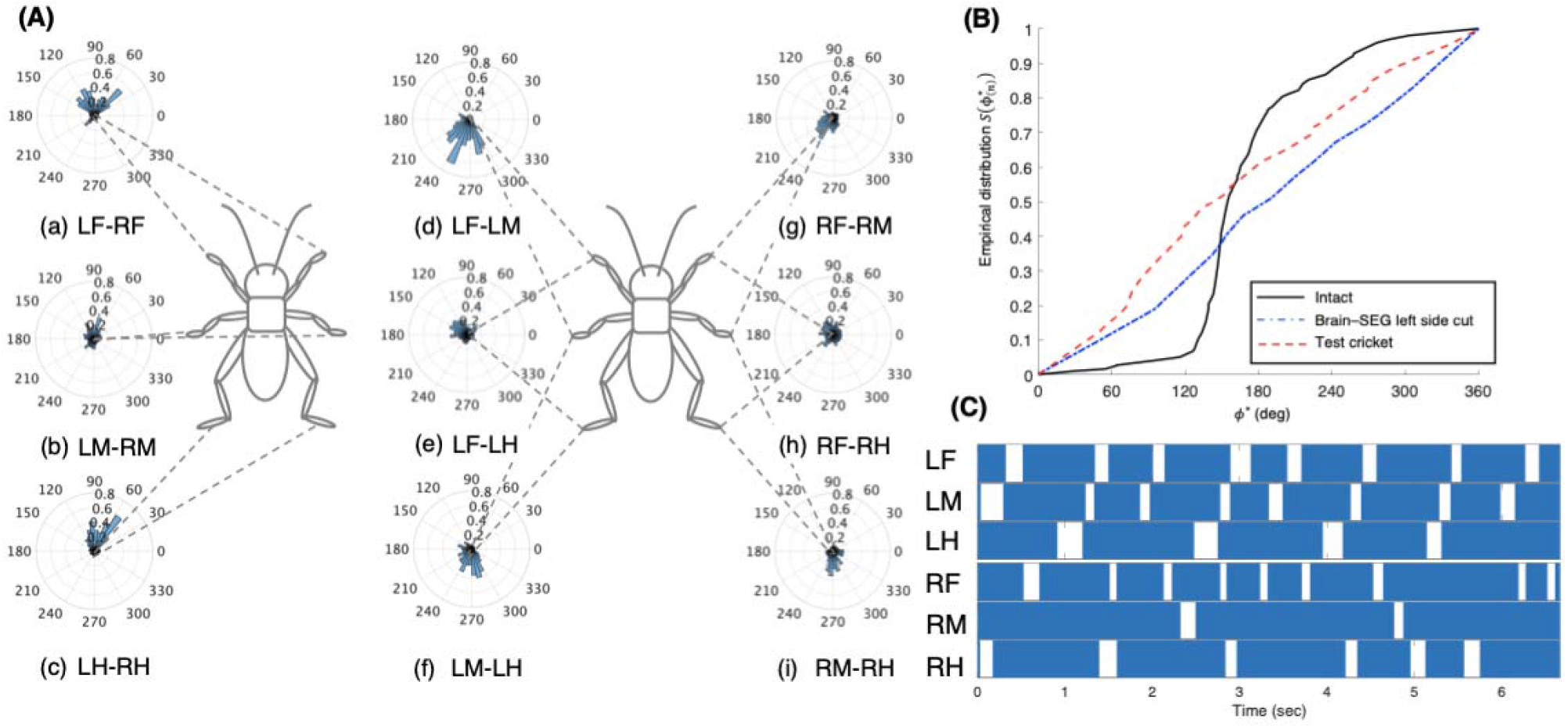
Walking gait patterns of crickets in which left sides of nerve connectives between brain and SEG, and between SEG and prothoracic ganglion were cut. **(A)** Polar histograms showing phase differences between two legs in test crickets (N = 3). The test crickets continued to turn in the right direction. The polar histograms demonstrate that the walking pattern was not a tripod gait, (a) to (i) Phase differences between pairs of legs. **(B)** Comparison of empirical distribution function of phase differences between left and right legs *Φ_LM–RM_*. The black line indicates the empirical distribution function of the intact crickets, the blue line indicates that of the test crickets, and the red line indicates that of the crickets in which the left side of the nerve connectives between the brain and SEG was cut (shown in Fig 3(**B**)). **(C)** Gait chart diagram of test cricket. The filled part indicates the duration of the power stroke period and the blank part indicates the duration of the recovery stroke. This demonstrates that the walking pattern was not a tripod in the test crickets. LF: left foreleg, RF: right foreleg, LM: left midleg, RM: right midleg, LH: left hindleg, and RH: right hindleg.

The behavior of crickets in which the left–side connective between the brain and SEG, and the right–side connective between the SEG and prothoracic ganglion were cut was the same as that of the crickets in which the left–side connectives between the brain and the SEG, and between the SEG and prothoracic ganglion were cut. Again, the test crickets did not walk straight forward, but continued to turn in the right direction (N = 3, Sup5 movie). The gaits of these test crickets did not exhibit a tripod pattern (Fig. 7). The polar histogram of the test crickets indicates that the phase differences between the left and right legs were not consistent (Fig. 7(A)). In the test crickets, the mean of the midleg phase difference *Φ_RM–RM_* was 87.5°, with a standard deviation *ν_LM–RM_* of 110°. The mean vector length *R_RM–RM_* was 0.16. The shape of the empirical distribution function of the midlegs of the test crickets was different from that of the intact crickets (Fig. 7(B)). The gait chart diagram of the test crickets demonstrates that the duration of the left leg movements appeared to be rhythmic, as in the intact crickets, but the right legs were not coordinated (Fig. 7(C)). When the right–side connective between the brain and SEG, and the left–side connective between the SEG and prothoracic ganglion were cut, the test crickets continued to turn left (N = 3).

**Figure 7.**
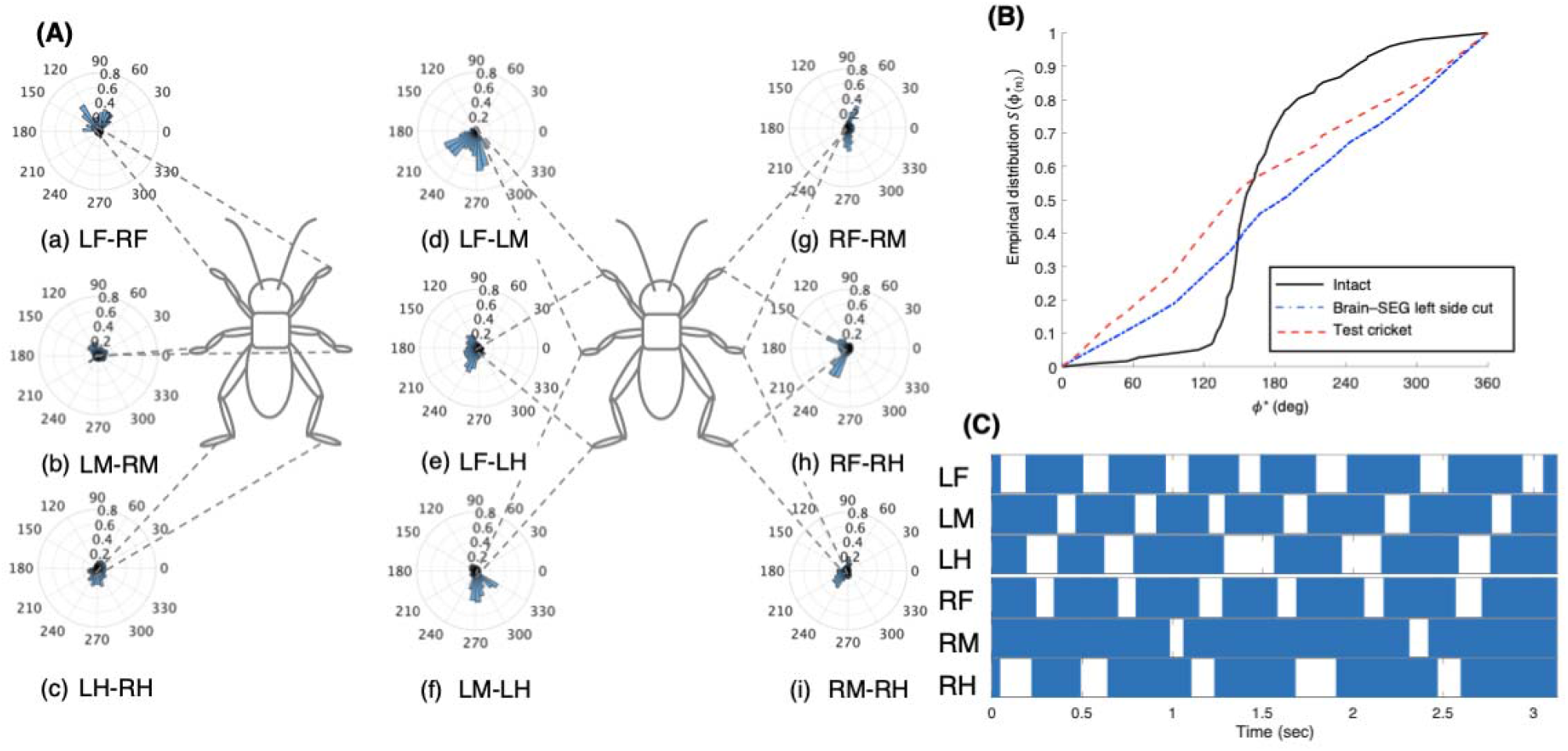
Walking gait patterns of crickets in which left side of nerve connectives between brain and SEG, and right side of connective between SEG and prothoracic ganglion were cut. **(A)** Polar histograms showing phase differences between two legs in test crickets (N = 3). The test crickets continued to turn in the right direction. The polar histograms demonstrate that the walking pattern was not a tripod gait, (a) to (i) Phase differences between pairs of legs. **(B)** Comparison of empirical distribution function of phase differences between left and right legs *Φ_LM–RM_* The black line indicates the empirical distribution function of the intact crickets, the blue line indicates that of the test crickets, and the red line indicates that of the crickets in which the left side of the nerve connectives between the brain and SEG was cut (shown in Fig 3(**B**)). **(C)** Gait chart diagram of test cricket. The filled part indicates the duration of the power stroke period and the blank part indicates the duration of the recovery stroke. This demonstrates that the walking pattern was not a tripod gait in the test crickets. LF: left foreleg, RF: right foreleg, LM: left midleg, RM: right midleg, LH: left hindleg, and RH: right hindleg.

### 3.3 Disconnection of connectives between metathoracic ganglion and abdominal ganglia

The crickets in which the pair of connectives between the metathoracic ganglion and first free abdominal ganglion was cut did not exhibit a tripod gait (Fig. 8(A), Sup6 movie). The phase differences between the left and right midlegs were not consistent (Fig. 8(A)(a)). In the test crickets, the mean of the mid-leg phase difference *Φ_RM–RM_* was 199°, with a standard deviation *ν_LM–RM_* of 108°. The mean vector length *R_RM–RM_* was 0.17. The shape of the empirical distribution function of the midlegs of the test crickets was far from that of the intact crickets (Fig. 8(B)). However, the walking gait pattern in the crickets in which the left–side connective between the metathoracic ganglion and first free abdominal ganglion was cut exhibited an ordinary tripod gait pattern (Fig. 9(A)). The polar histogram of the test crickets in which the left–side connective was cut indicates that the phase differences between the left and right legs occurred in an anti-phase manner (Fig. 9(A) (a)). In the test crickets, the mean of the midleg phase difference *Φ_RM–RM_* was 180°, with a standard deviation *ν_LM–RM_* of 55.7°. The mean vector length *R_RM–RM_* was 0.62. The shape of the empirical distribution function of the midlegs of the test crickets was similar to that of the intact crickets (Fig. 9(B)). Similarly, when the right–side connective between the metathoracic ganglion and third abdominal ganglion was cut, the test crickets exhibited a tripod gait walk, as in the intact crickets.

**Figure 8.**
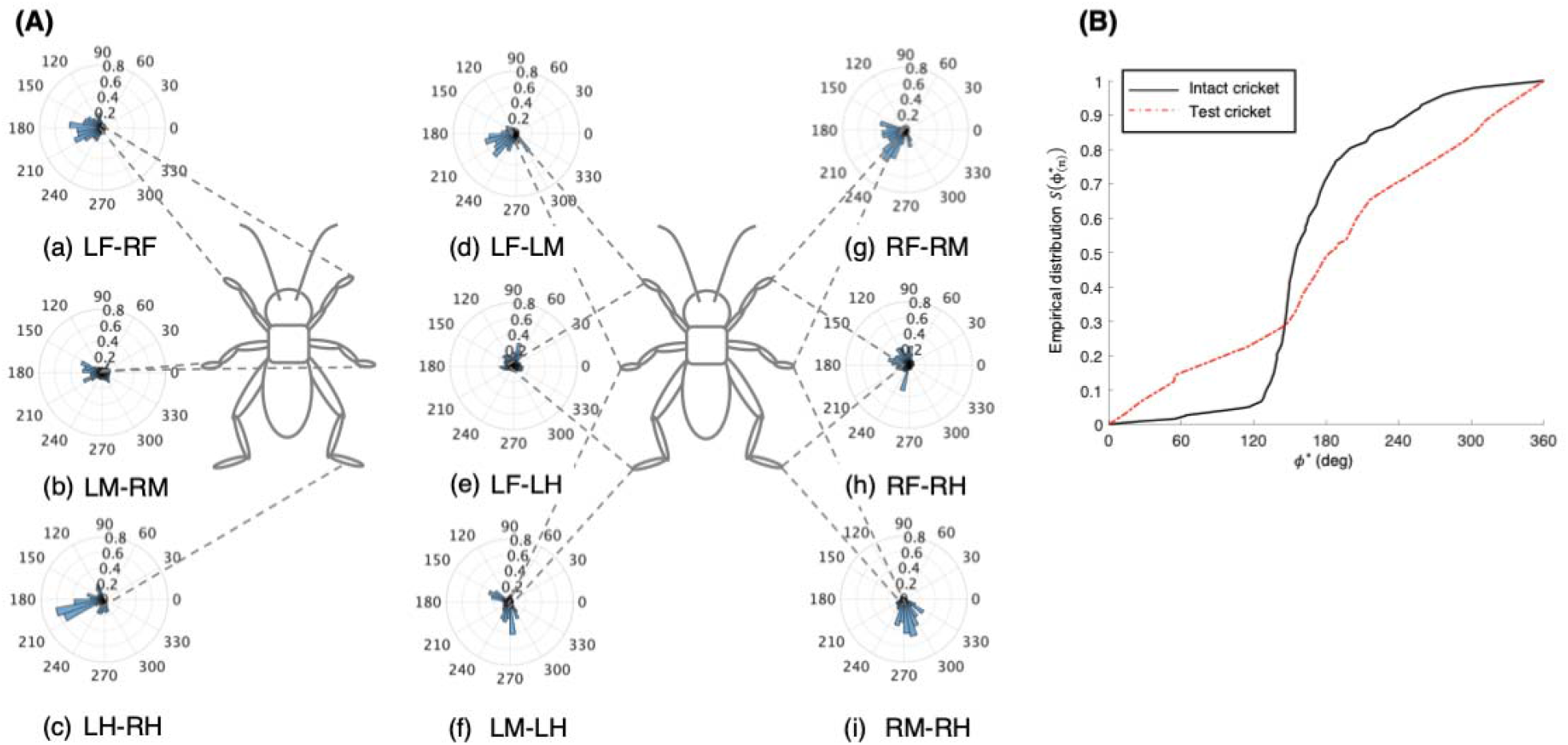
Walking gait patterns of crickets in which paired nerve connectives between metathoracic ganglion and abdominal ganglia were cut. **(A)** Polar histograms showing phase differences between two legs in test crickets (N = 5). The polar histograms demonstrate that the test crickets did not exhibit a tripod gait pattern on the treadmill, (a) to (i) Phase differences between pairs of legs. **(B)** Comparison of empirical distribution function of phase differences between left and right legs *Φ_RM–RM_*. The black line indicates the empirical distribution function of the intact crickets and the red line indicates that of the crickets in which the paired nerve connectives between the metathoracic ganglion and abdominal ganglia were cut. LF: left foreleg, RF: right foreleg, LM: left midleg, RM: right midleg, LH: left hindleg, RH: right hindleg.

**Figure 9.**
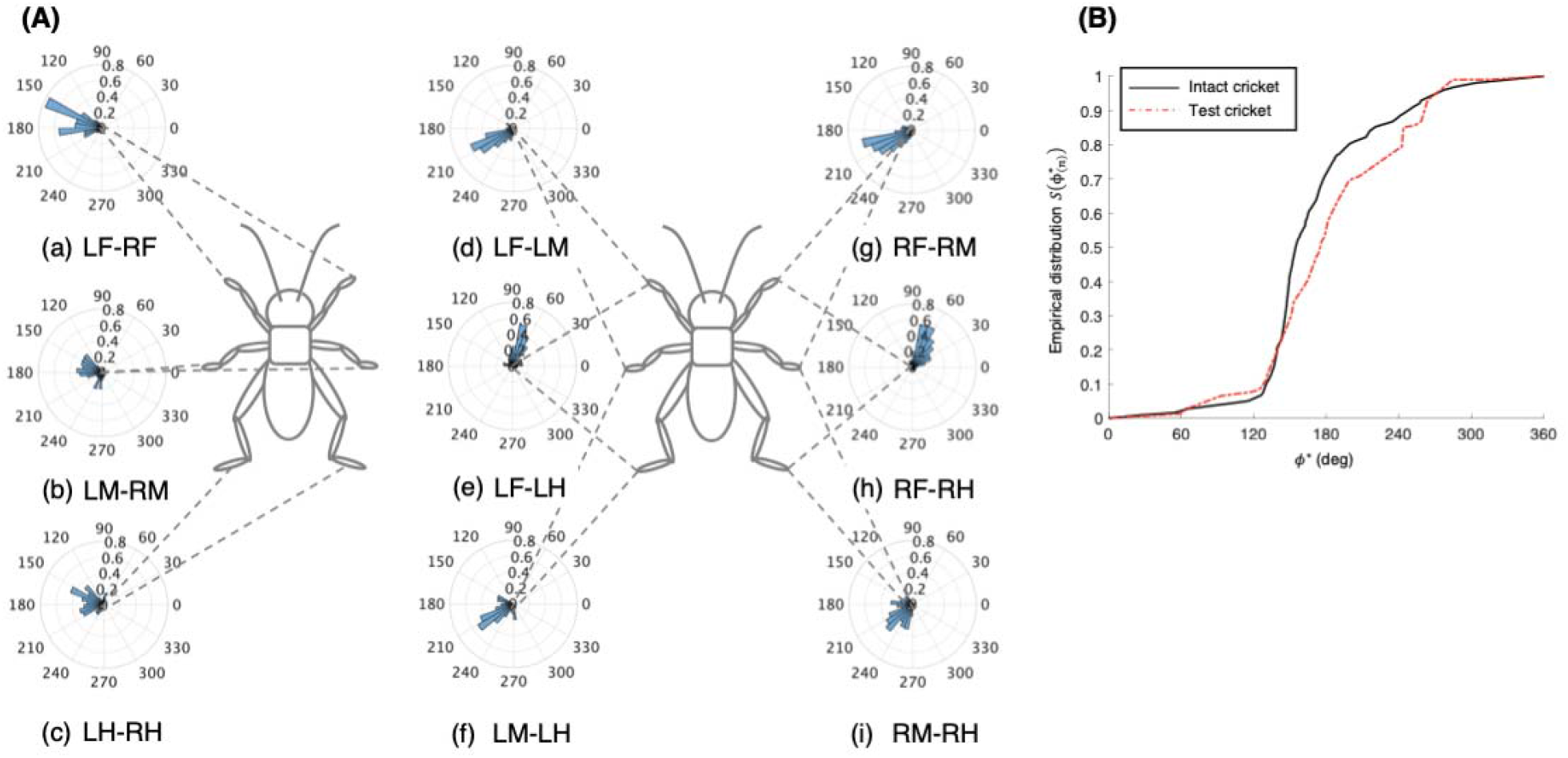
Walking gait patterns of crickets in which left side of nerve connectives between metathoracic ganglion and abdominal ganglia was cut. **(A)** Polar histograms showing phase differences between two legs in test crickets (N = 5). The polar histograms demonstrate that the test crickets exhibited a tripod gait pattern on the treadmill, (a) to (i) Phase differences between pairs of legs. **(B)** Comparison of empirical distribution function of phase differences between left and right legs *Φ_LM–RM_*. The black line indicates the empirical distribution function of the intact crickets and the red line indicates that of the crickets in which the left side of the nerve connectives between the metathoracic ganglion and abdominal ganglia was cut. LF: left foreleg, RF: right foreleg, LM: left midleg, RM: right midleg, LH: left hindleg, and RH: right hindleg.

## 4 Discussion

Crickets walk with a tripod gait pattern on a flat floor. Although the tripod gait is typical in insect walking, the gait patterns are not always fixed, but rather, change flexibly depending on the ground surface structure. The walking gait patterns may also vary if **the body structure is changed; for example**, owing to a **loss of legs** as a result of **an accident** (Full and Tu, 1991). To evaluate the changes in the gait patterns, gait chart diagrams of insects have been drawn in many previous studies (Wilson, 1966). A gait chart diagram has also been used to evaluate **the gait pattern of legged robots** (Owaki et al., 2017). Such a diagram expresses the movements of each leg and clearly indicates a snapshot of the position of each leg. Moreover, polar histograms describing gait patterns evaluate the phase differences of a given pair of legs during walking (Naniwa et al., 2020). One of the advantages of using a polar histogram is that it enables us to evaluate the gait patterns of not only individuals, but also a group of animals and legged robots. We investigated the effects of the loss of either the descending signals or ascending signals into the thoracic ganglia on regulating the cricket gait pattern.

### 4.1 Descending signals into thoracic ganglia to initiate walking

Although the crickets in which the paired connectives between the brain and SEG were cut could walk, they did not change their direction while walking on the treadmill. The polar histograms of the treated crickets demonstrated that their gait was very close to the typical tripod pattern of the intact crickets. As voluntary walking is initiated by descending signals originating in the brain (Kagaya and Takahata, 2011), the walking of the treated crickets was different from voluntary walking but could be initiated by receiving exteroceptive stimuli. The crickets responded to either tactile stimuli or air puffing on the cerci while walking. Crickets detect air currents using filiform hairs that are arranged on the surface of the cerci of the abdomen, and respond with rapid avoidance movement when they are deflected (Edwards and Palka, 1974). Information on air movements is processed and integrated in the terminal abdominal ganglion, and the signals are transferred to the thoracic ganglia to initiate avoidance walking (Mendenhall and Murphey, 1974) (Aonuma et al., 2008) (Yono and Aonuma, 2008). Furthermore, the ascending signals from the abdominal nervous system also contribute to the initiation of walking; for example, after–defecation walking (Naniwa et al., 2019). Thus, certain types of internal or external stimuli contribute to the initiation of walking in brainless crickets. It has been reported that the brain inhibits all reflex activities (Bethe, 1898). Neuronal signals for coordinating the leg movements are generated in the thoracic ganglia of insects. A decrease in the inhibition from the brain may have contributed to the treated crickets walking straight forward in this study.

One of the remarkable findings of this study is that the crickets in which one side of the connectives between the brain and SEG was cut exhibited walking, while it continued to turn in the opposite direction to that of the surgical cut of the connective (Fig. 3). This phenomenon appeared as though the inhibition from the brain to the cut-side pathway was abolished. The legs on the side of the cut connective moved more, which in turn pushed the body to the opposite side to continue turning. Movements of the opposite side could be introduced when they were bent. Therefore, the movements of the opposite side legs appeared to be caused by the local reflex. Movements of the legs in insects are detected by proprioceptive receptors (Tuthill and Wilson, 2016) such as the chordotonal organs (Hofmann et al., 1985) (Büschges, 1994), campaniform sensilla (Bässler, 1977), and hair plate (Pearson et al., 1976) (Wong and Pearson, 1976). Moreover, sensory afferents directly activate the extensor motor neurons of the trochanter and directly inhibit the flexor motor neurons in the cockroach (Pearson et al., 1976). Thus, bending the leg joints could activate the directory extensor motor neurons to extend the legs of the crickets. Therefore, our results suggest that inhibition of the brain contributes to the regulation of coordinated walking in crickets.

The descending signals from the SEG into the thoracic ganglia are important for initiating walking. The crickets in which the paired connectives between the SEG and prothoracic ganglion were cut did not walk, except after defecation, as reported for the behavior of the headless cricket (Naniwa et al., 2019). The motor neurons that activate the leg muscles originate in the thoracic ganglia. The rhythmic activities of neurons, known as CPGs, in the thoracic ganglia are thought to be closely linked to coordinated leg movements (Büschges et al., 1995) (Büschges, 1998) (Ritzmann and Büschges, 2007). The CPGs are modulated by the descending signals from the brain that initiate, maintain, modify, **and stop the motor outputs for walking** (Bidaye et al., 2017). The roles of the SEG are believed to modulate **the interactions between** the **sensory inputs from the legs and motor output** (Knebel et al., 2018a). Our behavior experiments confirmed the important role of the SEG in initiating walking.

Another significant finding in this study is that the crickets in which one side of the paired connective between the SEG and prothoracic ganglion was cut walked like the intact crickets (Figs. 5 and 9). Furthermore, the crickets in which one side of the connectives between the brain and SEG, and between either the ipsilateral or opposite side of the SEG and prothoracic ganglion were cut continued to turn in the opposite side to that of the connective cut between the brain and SEG (Figs. 6 and 7). This suggests that the descending signals from the SEG converge and are processed in the thoracic ganglia, and that the leg movements are regulated by the information from the SEG, even if it is only passed through one side of the connectives. Therefore, neurons may exist that integrate the information passed through the left and right pathways. Bilaterally symmetrical dorsal unpaired median (DUM) neurons have been identified in insects (locust: (Plotnikova, 1969); cockroach: (Crossman et al., 1971); and crickets (Clark, 1976)). Certain DUM neurons terminate in the leg muscles of cockroaches (Denburg and Barker, 1982) (Yoshitaka and Hiroshi, 1988). Moreover, the DUM neurons in the prothoracic ganglion contribute to walking regulation in crickets (Gras et al., 1990). Further investigation is required to clarify which neurons contribute to interlimb coordination in crickets.

### 4.2 Effect of ascending signals from abdominal nervous system on walking

In insects, the abdominal nervous system serves as the center for controlling avoidance behavior (Mendenhall and Murphey, 1974) (Tauber and Camhi, 1995) (Card, 2012), mating behavior (Killian et al., 2006), egg laying behavior (Sugawara and Loher, 1986), and defecation walking (Naniwa et al., 2019). These behaviors are closely linked to walking. Therefore, ascending signals from the abdominal ganglia into the thoracic ganglia may contribute to initiating and regulating walking in crickets. Furthermore, the descending signals that are modulated by the sensory feedback signals from the legs contribute to the modulation **of the coordinated walking gait** (Bidaye et al., 2017) (Knebel et al., 2018b). This study demonstrated that the ascending signals from the abdominal nervous system into the thoracic nervous system also contribute to coordinating walking gait patterns in crickets. The disconnection of the paired connectives between the metathoracic ganglion and first free abdominal ganglion prevented tripod gait walking in the crickets. However, the disconnection of one side of the connectives between the metathoracic ganglion and first free abdominal ganglion did not affect the expression of the tripod gait. Therefore, similar to the descending signals from the SEG into the thoracic ganglia, the ascending signals may be transferred into the bilateral neurons to be integrated and processed in the thoracic ganglia. Coordinated walking gait patterns are thought to be produced by the CPGs, descending central commands, and sensory feedback loops. This study demonstrated that the ascending signals from the abdominal nervous system also contribute to the generation of coordinated walking gait patterns in insects. Thus, it is necessary to investigate which types of neurons contribute to regulating the leg movements in crickets.

## Author contributions

KN and HA conceived and designed the experiment, performed the experiment, analyzed the data, and wrote the manuscript.

## Acknowledgments

We thank Professors K. Osuka and Y. Sugimoto (Osaka University) for their comments on this study.

